# Long-term experimental evolution decouples size and production costs in *Escherichia coli*

**DOI:** 10.1101/2021.09.28.462250

**Authors:** Dustin J. Marshall, Martino Malerba, Thomas Lines, Aysha L. Sezmis, Chowdhury M. Hasan, Richard E. Lenski, Michael J. McDonald

## Abstract

Body size covaries with population dynamics across life’s domains. Theory holds that metabolism imposes fundamental constraints on the coevolution of size and demography. However, studies of interspecific patterns are confounded by other factors that covary with size and demography, and experimental tests of the causal links remain elusive. Here we leverage a 60,000-generation experiment in which *Escherichia coli* populations evolved larger cells to examine intraspecific metabolic scaling and correlations with demographic parameters. Metabolic theory successfully predicted the relations among size, metabolism, and maximum population density, with strong support for Damuth’s law of energy equivalence in this experiment. In contrast, populations of larger cells grew faster than those of smaller cells, contradicting the fundamental assumption that costs of production should increase proportionately with size. The finding that the costs of production are substantially decoupled from size requires re-examining the evolutionary drivers and ecological consequences of biological size more generally.

## Introduction

The size of individual organisms drives widespread and repeated patterns across the tree of life^1-4^. For example, Damuth’s Rule holds that larger organisms have lower population densities than smaller organisms^5^. Similarly, populations of larger organisms grow slower than populations of smaller organisms^6^. Meanwhile, global warming and harvesting are causing declines in body size within many species, from phytoplankton to fish^7-10^. If body size and demography covary within species as they do across species, then human-induced changes in body size may have profound consequences for ecosystem function, particularly with regards to food security and the global carbon pump^11^. However, our ability to anticipate such changes is limited by the dearth of studies examining the within-species covariance of size, energy, and demography.

Metabolism has long been argued to provide the mechanistic link between size and demography because it governs the rate at which organisms transform energy into biological work and growth^4-6^. Larger species have higher absolute metabolic rates than smaller species, but lower metabolic rates relative to their mass. In formal terms, absolute metabolism scales hypoallometrically with body size with an exponent of *B*, whereas mass-specific metabolism scales at *B* – 1. The hypoallometric scaling of size and metabolism generates several predictions for how size should affect demography^12^.

First, because the ability to perform biological work per unit mass should scale with mass-specific metabolic rates, maximum rates of population growth (*r*) should also scale at *B* – 1 ^12,13^. For metazoans, *B* is typically ∼0.75; thus, *r* should scale around –0.25, which is strongly supported by interspecific comparisons^4^. This prediction has intuitive appeal: mouse populations can grow much faster than elephant populations.

Second, smaller species should attain higher population densities (*K*) than larger species, because their absolute *per capita* demands are lower. The resource requirements of organisms depend on their metabolism, so populations of larger species should cease growing at lower densities than those of smaller species^5^. However, larger organisms have lower mass-specific metabolic rates (in metazoans, at least), and so they require fewer resources per unit mass than smaller organisms. Accordingly, populations of larger organisms should have greater total masses at carrying capacity than populations of smaller organisms, with the expected scaling at 1 – *B* ^1^. This relation is known as the theory of energy equivalence^3^.

Finally, the maximum rate of population productivity (effectively the product of *r* and *K*) should scale with size at –1 when expressed as the rate of production of individuals, and so it should be size-independent (i.e., scaling exponent of 0) in terms of the rate of biomass production^2,12^. Together these three core predictions represent the canonical elements of how size, metabolism, and energy equivalence determine population growth and dynamics. Put simply, populations of larger organisms, with lower mass-specific metabolic rates, should grow more slowly, but eventually achieve higher biomasses, than populations of smaller organisms^4^. Nonetheless, there remains a fundamental disconnect between theory and evidence: most tests are based on among-species comparisons, making it difficult to attribute metabolism as the underlying driver of such patterns.

Although metabolic theory successfully predicts variation in demography across the domains of life, these predictions often falter when applied to narrower groups of taxa^2,12,14,15^. Various explanations have been offered for these discrepancies, but a key difficulty lies in inferring causality with respect to size differences across species. Mice differ from elephants in ways other than size, but metabolic theories about the relation between size and demography ignore these differences, treating them as an error term that is uncorrelated with size. We know, however, that many other traits covary with size (e.g., lifespan generally increases with size), and these traits also affect population dynamics^4,13^. Interspecific comparisons of individual size and population dynamics therefore confound other species-specific traits that influence demographic variables. Consequently, it remains unclear whether size, energy, and population dynamics are invariably related as supposed by the canonical scaling theory. Meanwhile, our capacity to predict the consequences of human-mediated impacts on the size of organisms depends on understanding the causal links between these factors within species.

Intraspecific tests of the relation between body size and demography are challenging. Comparisons among individuals of the same species suffer from limited power because they compare a narrower range of sizes than comparisons across species. Intraspecific comparisons of individuals at different ontogenetic stages can span a greater size range, but this approach also introduces confounding factors and cannot be extended to demographic parameters that must integrate across all ontogenetic stages. Ideally, a species that varies significantly in size across populations, and that allows the direct parameterisation of population dynamical models, would provide valuable evidence of how intraspecific variation in size and metabolic rates affects demography. However, such tests are rare^2,11^, and they have typically relied on temperature manipulations or strong artificial selection to generate differences.

Here we analyse the relations among organismal size, metabolism, and demography in 12 populations of *Escherichia coli* that have evolved and diverged from a common ancestor in the Long-Term Evolution Experiment (LTEE) over a period of 60,000 generations^16^. The LTEE populations have been thoroughly characterised, including by competitive fitness assays as well as whole-genome and whole-population sequencing^17,18^. Over the duration of the LTEE, each population has steadily increased in fitness, while accumulating many mutations. In this study, we measure their population dynamics, metabolism, and cell size to determine how these factors covary with each other, thereby allowing us to test whether they conform to predictions based on standard metabolic theory. In particular, we examine population growth rates and yields and find that the evolution of larger cell sizes has led to ‘Pareto improvements’ whereby growth rate has increased but not at the expense of yield^19,20^.

## Results

We examined two clones from each of the 12 LTEE populations at the 10,000 and 60,000 generation time points. We excluded the 60,000-generation clones from one population (Ara–3) that evolved the ability to consume citrate^21,22^, which is present in the medium as a chelating agent, because it gives cells access to an additional resource that confounds the relation between metabolism and demography that we seek to understand. In all analyses, we treat the average value of the two clones from the same population and generation as a single sample. We also include the two ancestral strains, REL606 and REL607, each of which was used to found six populations, and which differ by a genetic marker used in competition assays^16,18,23^. Thus, our analyses include a total of 25 samples (2 ancestors, 12 populations at 10,000 generations, and 11 populations at 60,000 generations).

Previous studies reported large increases in cell volume in the first 50,000 generations of the LTEE^21,22^. Our measurements confirm the large increases in cell size and show that they have continued to increase, from an average of 0.239 fL (i.e., µm^3^) for the ancestors to an average of 0.670 fL for the 60,000-generation samples (Figure 1). The 12 lineages followed different size trajectories, but they all show the same trend of increasing size.

**Figure 1.**
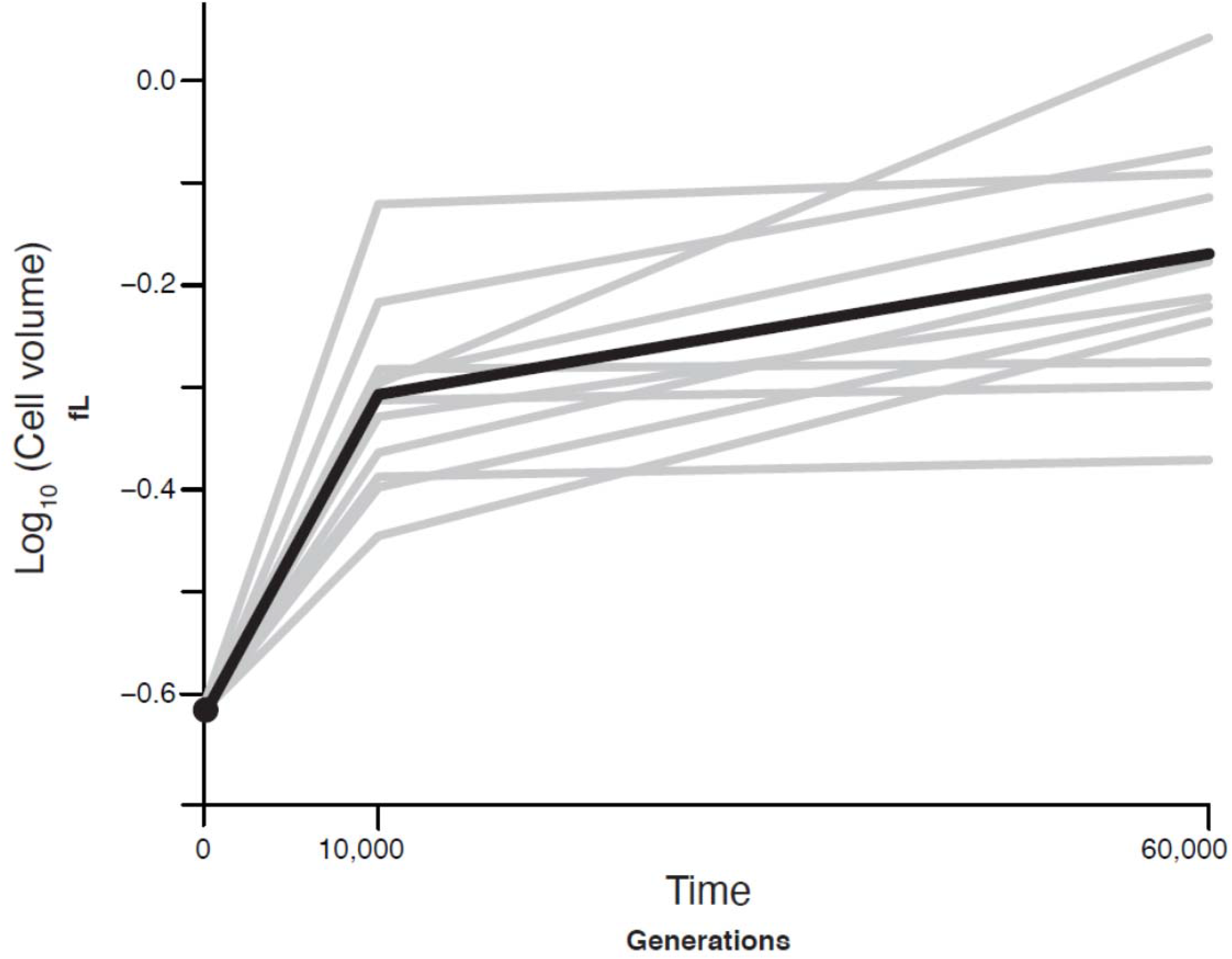
Trajectories of cell size in *E. coli* populations across 60,000 generations of evolution. Black line shows the mean trajectory of all populations; grey lines show the 12 independent populations.

We quantified metabolism by measuring oxygen consumption at three initial cell densities, achieved by differentially diluting samples. The concentration of the limiting resource, glucose, was the same for all three initial densities, and it was insufficient to support one population doubling even at the lowest initial density. As a result, the glucose was depleted over the course of our measurements of oxygen consumption, leading to a transition into stationary phase and concomitant decline in the *per capita* respiration rates at the higher initial densities. At all three initial cell densities, metabolism scaled with average cell size (volume) sub-linearly, and the scaling relation was consistent across the three densities (Density x log[Cell size]: F_2,69_ = 0.082, P = 0.921; Density: F_2,71_ = 97.06, P < 0.0001; log[Cell size]: F_1,71_ 45.99, P < 0.0001; Figure 2). The estimated scaling exponent for the metabolic rate, *B*, is 0.38, which differs significantly from interspecies comparisons^24^ that have estimated the scaling exponent to be >1, and from theoretical expectations based on surface-area-to-volume ratios of ∼0.67 – 1 (depending on cell shape). With our estimate of the intraspecific metabolic scaling exponent, we can then use standard metabolic theory to predict how population growth rates and maximum population size should scale with cell size (Table 1).

**Table 1.** Summary of predicted and observed scaling of population parameters based on metabolic scaling theory. We estimated the metabolic scaling exponent, *B*, as 0.38 (Figure 2). We show predictions (including confidence intervals in parentheses) based on standard theory, whereby production costs are assumed to scale perfectly with size (*C* = 1); when production costs are assumed to be size invariant (*C* = 0); and when production costs scale weakly with size (*C* = 0.11). The C value of 0.11 was calculated based on the scaling observed for the intrinsic rate of increase, *r*.

| Parameter | Definition | General theory | Prediction if $C = 1$ | Prediction if $C = 0$ | Prediction if $C = 0.11$ | Observed scaling |
| --- | --- | --- | --- | --- | --- | --- |
| $r$ | Intrinsic rate of increase | $M^{B-C}$ | -0.62 (-0.73:-0.51) | 0.38 (0.27:0.49) | 0.27 (0.16:0.38) | 0.27 (0.20:0.34) |
| $Max_{cells}$ | Maximum cell density | $M^{-B}$ | -0.38 (-0.49:-0.27) | -0.38 (-0.49:-0.27) | -0.38 (-0.49:-0.27) | -0.45 (-0.54:-0.37) |
| $Max_{biovolume}$ | Maximum population biovolume | $M^{1-B}$ | 0.62 (0.51:0.73) | 0.62 (0.51:0.73) | 0.62 (0.51:0.73) | 0.55 (0.46:0.63) |
| $Biovolume\ productivity$ | Maximum productivity | $M^{(1-B)} \times M^{(B-C)} = M^{1-C}$ | 0 | 1 | 0.89 | 0.81 (0.72:0.91) |

**Figure 2.**
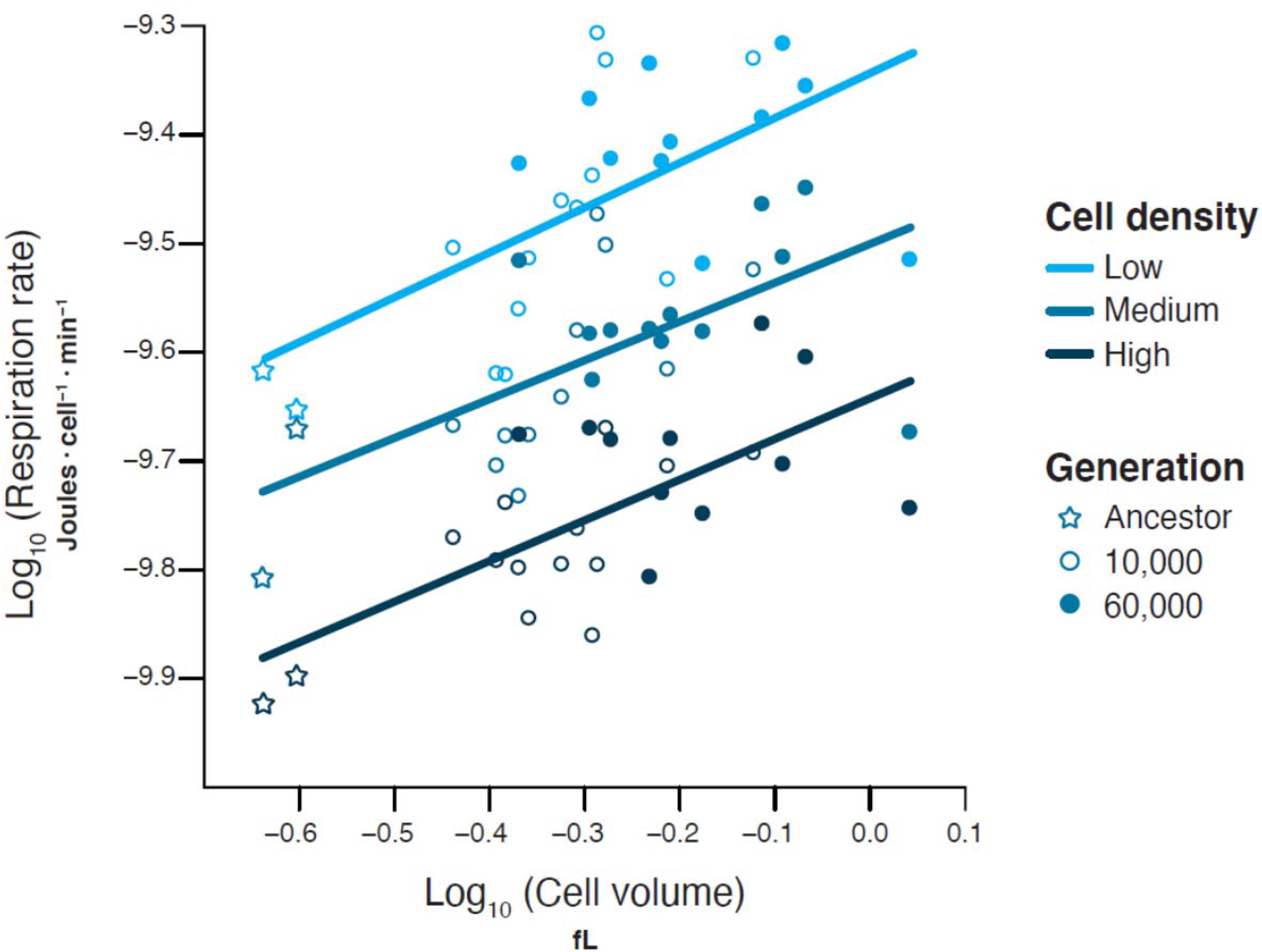
Scaling relation between average cell volume and *per capita* metabolic rate. The relation was examined across three different total biomasses, achieved by varying the initial cell density (shown by different colours). The limiting glucose concentration was the same for all three treatments; the glucose was thus depleted faster at the higher initial cell densities, leading to lower *per capita* metabolic rates at the higher densities. Each point shows the mean value for a sample at the generation indicated by the different symbols. The resulting overall estimate of the metabolic scaling exponent, *B*, is 0.38 and constant across densities.

We measured population growth over 24 h for the 25 samples, each at three different resource levels achieved by varying the concentration of glucose in the medium, and with replication of the growth curves at each concentration. Populations grew slightly faster at the higher glucose concentrations (Figure 3a). However, the scaling of the maximum growth rate, *r*, was consistent across glucose levels (Glucose x log[Cell size]: F_2,69_ = 0.113, P = 0.893). The scaling exponent of the growth rate was 0.27, which differs significantly from both zero and the exponent (–0.63) predicted by the canonical theory (Table 1). Instead, the scaling of the growth rate is much closer to that of the metabolic scaling (0.38 versus 0.27).

**Figure 3.**
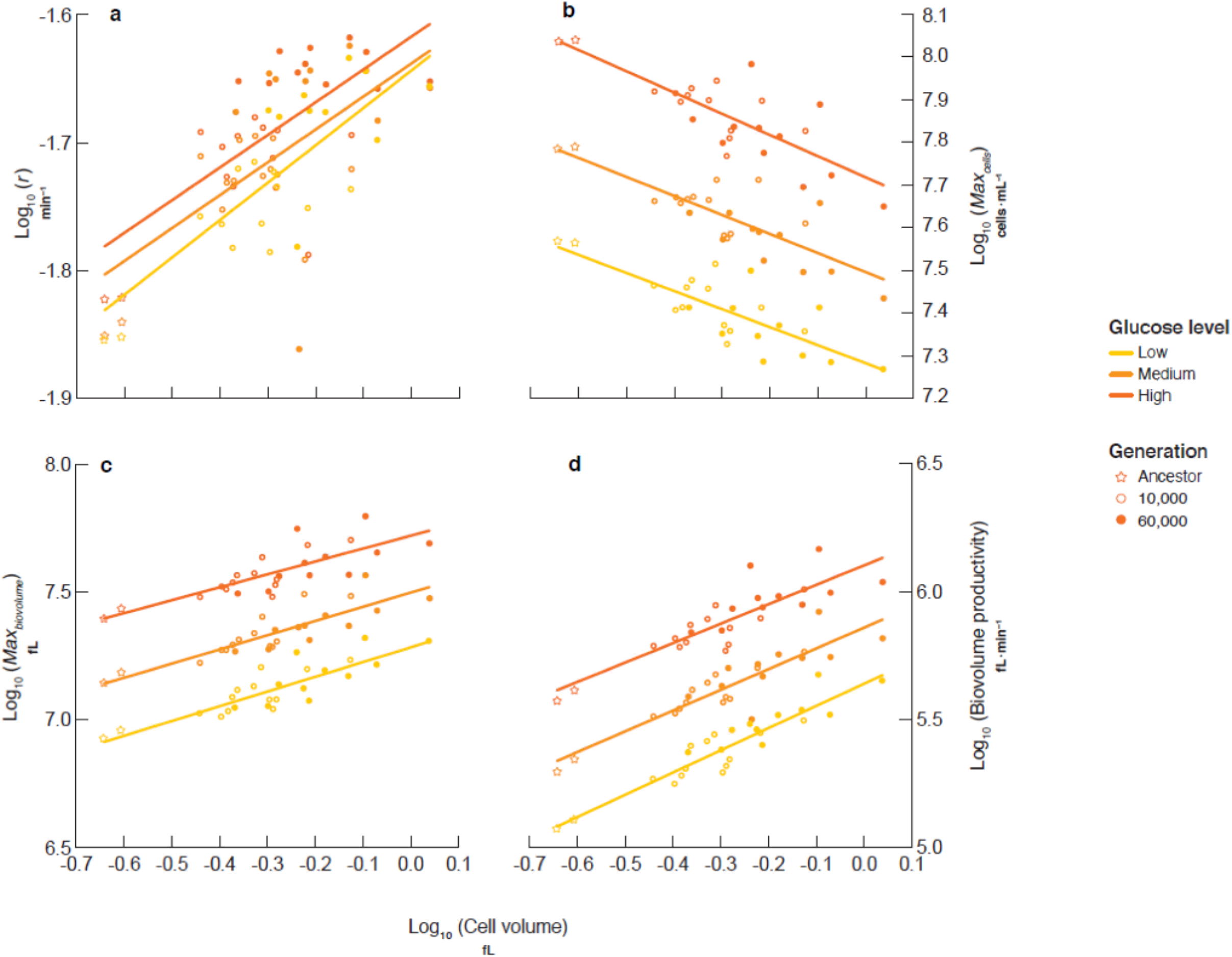
Scaling relations between average cell volume and a) intrinsic rate of population growth; b) maximum population density in terms of cell number; c) maximum population density in terms of total biovolume; and d) maximum rate of biovolume productivity. Different colours represent different glucose levels, with the lowest level equal to the concentration used in the LTEE. Each point shows the mean value for a sample at the generation indicated by the different symbols.

The maximum yield in terms of cell density (*Max*_*cells*_) showed a negative scaling relation with cell volume, with an exponent of –0.45 (Figure 3b), and the confidence interval overlaps the prediction of –0.38 from theory (Table 1). The correlation between cell size and maximum cell density was strong; a model including glucose level and cell size explained 96% of the variation in maximum cell density. The maximum biovolume yield (*Max*_*biovolume*_) scaled positively with cell size with an exponent of 0.55 (Figure 3c), again in reasonable agreement with the theoretical expectation of 0.64 (Table 1). As expected, populations achieved higher biovolumes at higher glucose levels, but the scaling relation was consistent across the three glucose levels (glucose: F_2,71_ = 437.32, P <0.0001; glucose x log[cell size]: F_2,69_ = 0.257, P = 0.774; Figure 3c).

Maximum productivity, expressed as the maximum rate of biovolume increase, increased with cell size, with an estimated exponent of 0.81 (Table 1). This estimate differs greatly, and significantly, from the canonical expectation of zero (Productivity_biovolume_: F_1,71_ = 301.5, P < 0.001). Productivity increased at higher glucose levels (glucose: F_2,71_ = 410.5, P <0.0001; Figure 3d), with no significant interaction between cell size and glucose levels (F_2,69_ = 0.447, P = 0.641).

Table 1 summarizes our results relative to theoretical expectations. Maximum population size scaled almost exactly as metabolic theory would predict, regardless of whether it was measured in terms of cell number (Figure 3b) or total biovolume (Figure 3c). In contrast, productivity did not conform to the predictions made by the canonical metabolic theory, whether it was measured as the rate of population increase (Figure 3a) or the maximum biovolume productivity (Figure 3d). Instead, both productivity exponents were much higher than the canonical theory would predict, by values of 0.89 and 0.81, respectively.

## Discussion

The LTEE provides a unique opportunity to study the covariance between size, metabolism. and demography within a species. Damuth’s law of energy equivalence successfully predicted the coevolution of cell size with maximum cell density^5^. However, our results indicate that a fundamental assumption about how the growth and productivity of populations should scale with metabolism and cell size lacks generality and therefore requires modification. Our study also indicates the need for more within-species tests of metabolic theory, because evolution can evidently lead to Pareto improvements in key size-related parameters—leading to trade-ups, rather than trade-offs—that are not anticipated from interspecific comparisons.

### Larger cells have relatively lower metabolic rates than smaller cells

Metabolic scaling in these *E. coli* populations is remarkably low, with an exponent of ∼0.38. Among-species comparisons of metabolic rate in bacteria have usually reported hyperallometric scaling (*B* > 1), whereby larger cells have disproportionately higher metabolic rates^24,25^. Instead, we find that the larger cells from later generations have much lower mass-specific metabolic rates than smaller cells, such that a 3-fold change in size results in only a 1.5-fold increase in metabolism.

There are several potential explanations for the low scaling exponent that we observe in this experiment relative to interspecific comparisons. First, it could be that within-species metabolic scaling is generally shallower than interspecific scaling in bacteria; to date, there are too few studies that have measured within-species scaling to compare them. In other taxa, metabolic scaling sometimes differs depending on whether it is estimated within or among species^26,27^. Theory predicts that, all else equal, the physics of resource limitation in slow-moving fluids should result in metabolic scaling exponents of about 0.33 ^28^, which is close to our estimate. The cytoplasm of bacterial cells is viscous and densely packed with DNA and other macromolecules^21,22^. It could also be that physical constraints on scaling are more restrictive within-than among-species, where cell shape may change along with cell size^28^. It should be noted, however, that the aspect ratio (length/width) also varies significantly among the *E. coli* lineages in this study^22^.

Second, the fine-tuning of gene regulation and physiological process may have led to the low metabolic scaling exponents seen in the LTEE. DeLong et al. suggested that hyperallometric metabolic scaling in bacteria emerges from the effect of genome size on metabolic rate^24^. Larger cells typically have larger genomes, and more genes and gene products might drive higher metabolic rates^29^. Although the average haploid genome length has declined slightly during the LTEE owing to some gene deletions^23^, rapidly growing bacterial cells typically have multiple copies of their chromosome. Therefore, the faster-growing and larger-sized evolved bacteria have more total DNA per cell, even if their genome length is slightly smaller. Among prokaryotes, genome length scales with cell size with an exponent of 0.35 ^24^, which is close to the 0.38 metabolic exponent we observed (Figure 2, Table 1). The bacteria in the LTEE have evolved substantial changes in gene expression and regulation^30-32^. These changes have reduced the expression of functions that are no longer useful in the LTEE’s simple conditions, while optimizing the expression of the functions that are needed in this predictable environment^33^.

### Metabolic theory predicts maximum population size

We found strong support for the energy equivalence rule across a range of resource levels^5^. Because the mass-specific metabolic rates of larger cells were so low, the maximum biovolume yields were much higher in the evolved samples than in the ancestors (Figure 3c). However, the total metabolic demands of these two groups were similar (∼4.5 × 10^−3^ J). Thus, the larger cells are metabolically more efficient and attain higher population biomass than smaller cells for a given amount of resource. This result conforms with other LTEE studies that found that the evolved cells are larger, more efficient, and attain higher maximum biomass yields than the ancestors^20,21^. It seems that metabolic rate can be an excellent predictor of the limits to population biomass, both among^4^ and within species^11^. In contrast, longstanding metabolic theories, based on standard assumptions, failed to predict how individual size and metabolism would impact population growth rates and maximum productivity.

### Metabolic theory and population growth rates

The *E. coli* samples in this study defy theoretical predictions based on standard assumptions about how individual size should affect rates of population growth and production. Despite having lower mass-specific metabolic rates, the larger evolved cells had higher intrinsic rates of increase (*r*) than the smaller ancestral cells. Larger cells should require more materials and energy to produce, but relative to their volume, they should also have less capacity to power this work relative to smaller cells. Nonetheless, our study, other studies of the LTEE populations, and indeed studies on *E. coli* more generally find that faster growing cells are larger than cells growing more slowly^20,21,34-36^. This positive correlation between size and growth rate contradicts standard theory, as well as intuition.

Standard theory predicts that population growth rate should scale with the mass-specific metabolic rate (i.e., M^B–1^) ^12,13,37^. This theory works well for among-species comparisons: in multicellular eukaryotes, both mass-specific metabolic rate and population growth rate scale at ∼M^−0.25 1,4^; and in prokaryotes, both rates scale at ∼1 ^24,38^. However, in the LTEE, population growth rate scales at 0.27, an exponent that is 0.89 higher than expected given the mass-specific scaling of –0.62. In fact, the population growth rate exponent is much closer to the *per capita* metabolic exponent of 0.38 than to the mass-specific exponent of –0.62. Why do these bacteria show positive scaling of both *per capita* metabolism and population growth rate with individual size, contradicting expectations based on the standard theory?

### Metabolic theory and the costs of biological production

A crucial, but often overlooked, assumption of standard metabolic theory is that the energy required to produce a new individual is directly proportional to its mass^6^. This assumption seems reasonable and intuitive at first glance, but in fact there is little empirical evidence to support it and, in the case of the LTEE, some evidence against it. The total cost of producing a cell is the sum of the energy consumed between cell divisions (sometimes called maintenance costs) and the energy used to build the new cell itself^38^. Neither component is likely to scale directly with cell volume, for several reasons.

First, it has been estimated that about half of the energy required by *E. coli* is used to maintain ion gradients across the cell membranes^39^. Larger cells have smaller surface area relative to mass, and so they should have relatively lower maintenance costs than smaller cells. Consistent with this reasoning, total metabolism scales hypoallometrically with cell volume in the LTEE. Second, large cells often have different stoichiometry from small cells. Both among and within taxa, large cells tend to have relatively lower carbon content than small ones^40^. In the LTEE specifically, size and carbon density do not scale proportionately, and the stoichiometry of cells has evolved over time^21,34^. In this light, the assumption of equal costs per unit volume of building smaller and larger cells is violated. Finally, large cells are relatively cheaper to produce than small cells in terms of genome replication. In the LTEE, the larger evolved cells have slightly smaller genomes than the smaller ancestral cells^23^, so that the relative, and even absolute, costs of genome replication are lower for the larger cells. Of likely greater importance, the evolved cells have undergone substantial fine-tuning of their gene-regulatory networks to the LTEE environment, thus reducing the costly expression of unneeded transcripts and proteins^30,31,33^.

### Relaxing the strict proportionality of production costs to cell size

Taken together, our results imply that larger cells are cheaper to maintain and build per unit volume, such that the scaling of the total cost of production is far less than proportional to cell size. If the assumption of proportional cost is relaxed, then the paradox of larger cells having higher growth rates may be resolved. Instead of assuming that the costs of production scale with individual cell size with an exponent of 1, we can explore a range of possible scaling exponents and compare the resulting predictions with our observations. To that end, here is the generalised formula relating cell size to population growth rate:

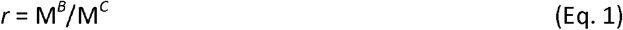

where *B* is the exponent linking cell mass to metabolic rate, and *C* is the exponent linking total production costs (both maintenance and building) to mass. When the costs are assumed to be directly proportional to size (i.e., *C* = 1), we recover the prediction of classic metabolic theory^13^:

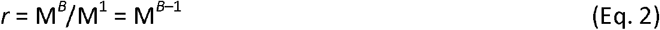

At the other extreme, the costs of production are size invariant (i.e., *C* = 0). That is, the total costs of producing smaller and larger cells are the same, and theory instead predicts:

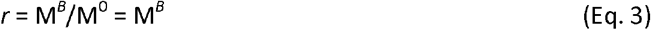

Of course, any value of *C* is possible in this more general framework. In the case of the LTEE strains, we find that *r* scales at 0.27, which implies that *C* = 0.11 (i.e., 0.38 - 0.27 = 0.11). In other words, the costs of production increase only weakly with cell size. Specifically, the cells from generation 60,000 are, on average, over twice the volume of their ancestors (Figure 1), but they cost only ∼10% more to produce than the small ancestral cells. If we now set the exponent that links production cost (*C*) to mass at 0.11, then we predict the scaling exponent for the maximum rate of biovolume production seen in our experiments (Table 1). In other words, if we assume the *per capita* cost of producing the larger evolved cells is only slightly more than the cost of the smaller ancestral cells, then we can reconcile our other observations with the classic theoretical predictions.

A recent study of the single-celled eukaryote *Dunaliella tertiolecta* also found improvements in both population growth rate and yield as cells evolved to be larger^2^. These improvements were associated with the evolution of significant genomic streamlining^41^, which likely decoupled some production costs from cell size. Thus, it seems that the trade-offs between size and rates of production that seem almost invariant in comparisons among species can be circumvented within species when other traits that affect metabolic costs also coevolve.

In conclusion, our results demonstrate the importance of examining the scaling of size, metabolism, and population dynamics within species, as well as across species, because these comparisons may differ quantitatively and even qualitatively. Such differences occur even though the explanations for these patterns at both scales involve the same underlying metabolic processes. Given the importance of the scaling of production costs to organismal size in driving our expectations of how size affects population growth and productivity^4^, this issue has received far too little empirical attention. We recommend, therefore, that future studies should examine production costs as a function of size, both within and among species.

## Acknowledgements

We thank Craig White and Nkrumah Grant for valuable discussions. This work was supported, in part, by the US National Science Foundation (grant DEB-1951307 to R.E.L.) and an ARC Future Fellowship (grant no. FT170100441) to MJM.

## Methods

### Experimental Overview

We measured average cell volumes for 48 *E. coli* clones (2 ancestral strains, 2 clones isolated from each of the 12 LTEE populations at 10,000 generations, and 2 clones from 11 populations at 60,000 generations; Table S1). We excluded from our analyses one population at 60,000 generations because it evolved the ability to use citrate as an additional source of carbon and energy in the LTEE environment. We measured metabolic rates of the same 48 clones at 3 initial cell densities. We monitored the population growth of the same 48 clones at each of 3 resource levels, to which we fit growth curves. We averaged the estimates of cell size, metabolic rate, and population growth parameters for the two evolved clones from the same population and generation. In all analyses, we treat the average value of the two evolved clones as a single sample. We also include the two ancestral strains, each of which founded six of the LTEE populations. Thus, our statistical analyses reflect a total of 25 samples (2 ancestors, 12 populations at 10,000 generations, and 11 populations at 60,000 generations) for each assay and, when relevant, for each treatment.

### Evolution Experiment, Strains, and Media

The *E. coli* long-term evolution experiment (LTEE) started in 1988^16^ and has continued since. Twelve 50-ml flasks containing 10 ml of DM25 medium (see below) were seeded with either the “arabinose-negative” ancestral strain REL606 (populations Ara–1 to Ara–6) or the “arabinose-positive” ancestor REL607 (Ara+1 to Ara+6). The Ara marker causes cells to produce either red (Ara^−^) or white (Ara^+^) colonies on tetrazolium-arabinose indicator plates, and it serves to differentiate competitors during relative fitness assays. The Ara marker is selectively neutral in the LTEE conditions^42^. The 12 populations are propagated daily with 100-fold dilutions at 37°C while shaking at 120 rpm for mixing and aeration. The dilutions and regrowth allow log_2_ 100 ≅ 6.6 cell generations per day. The stationary-phase (i.e., end of day) population density is ∼5∼10^7^ cells/mL for the ancestral strains,^16^. In 11 populations, the stationary-phase density declined as the individual cells became larger; in the case of population Ara–3, however, the cell density increased several-fold after cells evolved the new capacity to use the citrate in DM25 as an additional source of carbon and energy^43^. Samples (including whole populations and isolated clones) are periodically stored with glycerol (as a cryoprotectant) at –80°C, where the cells remain viable and available for further analyses.

Our analyses used the two ancestors, plus two clones sampled from each population at 10,000 and 60,000 generations (except for Ara–3 at 60,000 generations, which we excluded owing to its access to citrate as an additional substrate for growth). The isolation of the 10,000-generation clones was described previously^23^. For this study, we plated each 60,000-generation population sample on Lysogeny Broth (LB) agar and picked two clones at random, which we then stored as glycerol stocks.

The culture medium used in the LTEE and in this study is Davis Mingioli (DM) minimal medium [7 g/L potassium phosphate (dibasic trihydrate), 2 g/L potassium phosphate (monobasic anhydrous), 1 g/L ammonium sulfate, 0.5 g/L disodium citrate, 1 mL/L 10% magnesium sulfate, and 1 mL/L 0.2% thiamine (vitamin B1)] supplemented with a specified amount of glucose^16^. The concentration of glucose added to the medium is indicated by a suffix (e.g., DM25 has 25 mg/L glucose). MG agar plates were used for counting colonies; in addition to the ingredients of DM media, it contains 4g/L of glucose and 16g/L agar. LB broth [NaCl (10 g/L), tryptone (10 g/L), and yeast extract (5 g/L)] was used for the initial recovery of bacteria from thawed glycerol stocks for the haemocytometer count assay; LB plates were made by adding 20 g/L agar.

### Population Growth Measurements

Each clone was revived from the frozen stocks and then grown in 3 mL of DM25 at 37°C with orbital shaking for 24 h to acclimate the bacteria to that medium. The next day, we measured the optical density (OD) of each culture, and the density was normalised to match the culture with the lowest OD. The resulting cultures were diluted 100-fold into 96-well microplates containing DM25, DM50, or DM100 media. Each clone was replicated 4 times in each medium, for a total of 600 growth curves (50 clones x 3 media x 4 replicates, including the two clones from population Ara–3 at generation 60,000 that were subsequently excluded). The clones were randomly assigned to wells for each medium over 20 microplates to minimize position effects. We measured OD at 600 nm wavelength every 10 min for 24 h using an ELx808 Incubating Absorbance Microplate Reader (BioTek Instruments, USA) set to its maximum shaking speed and 37°C.

A complete description of the methods that we used to estimate demographic parameters is provided in Malerba et al.^44^. Briefly, OD serves as a proxy for population biomass, and we log_e_-transformed OD values to reduce heteroscedasticity. We then fit a four-parameter logistic-type sinusoidal growth model of the following form:

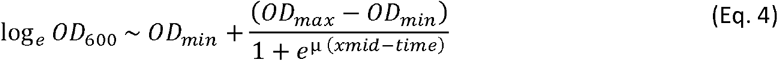

where *OD*_*min*_ is the minimum population biomass, *OD*_*max*_ is the maximum population biomass, *xmid* is the time to the inflection point, and μ quantifies the curve’s steepness. The following demographic parameters were extracted for each trajectory: the maximum predicted value for OD_600_ (*K*; unit: OD_600_); the maximum rate of biomass increase (*r*; unit: min^−1^); and the maximum rate of biomass production (unit: OD_600_ min^−1^).

### Metabolic Assays

We measured metabolic rates, based on oxygen consumption, as follows. The clones were revived from the frozen stocks by plating on LB agar. Single colonies were used to inoculate 2 mL of DM800 medium, and the cultures were incubated at 37°C with orbital shaking for 24 h. The next day, the cells were pelleted by centrifugation, washed with DM0 medium (i.e., DM without added glucose) to remove any residual glucose and extracellular by-products. The pellets were resuspended in 2 mL of DM0, and the cultures were then adjusted to OD_600_ values of 0.15, 0.3, and 0.6 and a final volume of 5 mL each using DM0.

Oxygen consumption was measured in a temperature-controlled room at 37°C using 4 × 24-channel PreSens Sensor Dish Reader (SDR; AS-1 Scientific Wellington, New Zealand), using methods adopted from Malerba et al.^27^. Before the experiment, the equipment was kept overnight in the 37°C room, and each SDR plate was calibrated using air-saturated (AS) DM800 medium (100% AS) and DM800 medium containing 2% sodium sulphite (0% AS). We monitored a total of 192 cultures that included the 2 ancestral and 48 evolved clones (including the two 60,000-generation clones from population Ara–3 that were later excluded) at each of the three initial cell densities, plus an additional 21 replicates of ancestral strain REL606 and 21 blanks without any cells. The additional ancestral replicates meant that each 24-well plate included this reference strain at all three cell densities, allowing us to detect possible plate-level anomalies; however, we encountered no such problems. The cultures were otherwise randomly distributed across two consecutive days of data collection. Each culture was carefully placed in a 5-mL vial to avoid creating any air pockets. At least two vials per plate were filled with sterile medium that served as blanks. Before starting the trials, all cultures were acclimated to 37°C for an hour. We added 0.4 µL of 10% glucose solution to each 5-mL sample prior to the start of the assays, which brought the glucose concentration to 8 mg/L (about one-third of the concentration in the standard LTEE medium, DM25). Moreover, even the lowest initial density (OD_600_ = 0.15) is higher than the final density the bacteria reach when they have depleted the glucose in DM25. Thus, the glucose supply was quickly exhausted during these metabolic assays, with the depletion occurring faster at the higher cell densities. This effect led to different estimates of metabolic rates across the three cell density treatments; however, the scaling exponent between cell volume and metabolic rate was unaffected by the treatment (Figure 2). The assays began after the SDR channels were fully loaded and the samples were well-mixed. The non-consumptive O_2_ sensors then monitored the oxygen in each vial every minute until it was consumed by the bacteria.

After the assays ended, the rate of change in oxygen saturation (VO2) was quantified from the linear part of each time-series (Figure S1). Energy rates were calculated with the following model:

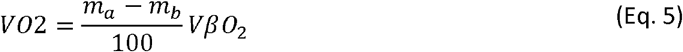

where *m*_*a*_ is the rate of change in each sample (% min^−1^), *m*_*b*_ is the mean rate of change for the blanks in,each plate (% min^−1^), *V* is the water volume (0.005 L), and (*βO*_2_ is the oxygen capacity of air-saturated water at 37°C and zero salinity (210 µmol O_2_ L^-1^). Finally, the rates were converted to energy units, assuming a calorific energy of 0.512 J (µmol O_2_)^−1^ from Malerba et al.^27^.

### Calibration Curves for OD and Cell Density

Calibration curves were required to convert oxygen consumption (VO2) and carrying capacity (*K*) from units of OD_600_ to units of cells per mL, and thereby express metabolism and productivity on a *per capita* basis. To this end, we measured cell densities, using two approaches. The first approach used a Neubauer Improved haemocytometer (Bright-line double ruled, Pacific Lab) to estimate cell densities for calculating *per capita* respiration rates. The bacteria were growing, at least briefly, during the respiration measurements, and therefore these calibrations used growing cultures. Clones were revived from glycerol stocks by inoculation into 1 mL LB medium and grown overnight. Cells were washed 3 times in 1 X PBS and then diluted 1000-fold in 3 mL of DM100 medium, where they grew at 37°C with orbital shaking for 24 h. The next day, the cultures were diluted 20-fold into 200 μL of DM400 medium in a 96-well microplate. We immediately measured an initial OD_600_ value for each well using the same ELx808 Incubating Absorbance Microplate Reader as for the population growth measurements. We then placed the plate in a Thermo Scientific plate shaker at 37°C and 750 rpm for 2 h. We recorded another set of OD_600_ readings, and then took a 20-μL sample from each well and diluted it to a final concentration of 5% formaldehyde to fix the cells. We returned the plate to the shaker at 37°C. Every hour, we recorded OD_600_ readings and took and fixed 20-μL samples for haemocytometer cell counts until 5 h had elapsed. Three to four replicate cultures were analysed for each clone, with a blinded set of clones used for measurements, which were conducted over 20 days. We rarely measured replicates from the same clone on a given day. Fixed cells were mixed by pipetting up and down, and we transferred 10 μL into the Neubauer chamber. We used a light microscope to count the cells. We ran a linear regression to convert optical density into haemocytometer-based cell densities for each sample, and the resulting values were used to convert oxygen consumption to *per capita* metabolic rates.

Maximum OD values typically occurred in our population-growth experiments when the cells had depleted the glucose and begun to enter stationary phase. Bacterial cells are smaller, on average, in stationary phase than while growing, including in the LTEE populations^22^. Therefore, the above calibrations could not be used to directly estimate maximum cell density (*Max*_*cells*_). Instead, we estimated the stationary-phase cell density after 24 h of each clone by plating cells on MG agar plates. Clones were revived from frozen stocks and grown in DM25. Aliquots of these cultures were distributed at random over multiple 96-well microplates to minimize position effects. After 24 h at 37°C on a plate shaker, each culture was diluted 1:100 in DM25, DM50, and DM100 (2 μL of culture in 200 μL of medium) and incubated again for 24 h on the plate shaker. These cultures were diluted 10,000-fold and spread on MG agar plates, and colonies were counted after incubating the plates for 24 h. We used these counts to calibrate stationary-phase cell densities based on colony-forming units at 24 h (N_CFU_) and cell densities inferred from OD values and haemocytometer counts of growing cells (N_OD_), which yielded the following equation: log_10_[N_CFU_] = 0.92 x log_10_[N_OD_] + 1.35 (see Figure S2). We then used this equation to estimate *Max*_*cells*_ as the cell density corresponding to the maximum OD reading (OD_max_) from each growth curve. The 60,000-generation sample from population Ara+3 appeared to be an outlier when calibrating the relation between cell numbers based on OD and CFU values (Figure S2). (Note: This outlier is not the Ara–3 sample that was excluded from all of our analyses because the cells can grow on citrate). We therefore recalculated all the scaling exponents in this work while excluding the outlier sample, but none of the values changed substantively (Table S2).

### Cell Size Measurements

We measured the mean cell volume for each clone in stationary phase using the side-scatter of a flow cytometer (Flow-Core, BD LSRII, BD Biosciences, San Jose, CA, USA); beads of four diameters (0.2, 0.5, 1, and 2 μm, Invitrogen by Thermo Fisher Scientific) served as standards. The clones were revived from frozen stocks and grown in DM25 at 37°C with orbital shaking for 24 h. The next day, these acclimated cells were diluted 100-fold in fresh DM25 medium in 96-well microplates. We had four replicates per clone, and the clones were randomly placed across four plates. The plates were incubated at 37°C and 750 rpm for another 24 h, at which time samples were taken for flow cytometry.

### Statistical Analyses

Metabolic rates and growth models were calculated using R^45^ and the packages nlme^46^, lme4^47^, and plyr^47^ for model fitting. ANCOVA and multiple regression models were performed to examine the scaling relations between average log-transformed cell volume and the various log-transformed metabolic and population dynamics metrics, respectively; initial cell density (in the case of metabolism) and glucose level (in the case of population dynamics) were additional covariates or fixed factors. In all cases, we calculated mean values across technical replicates for a given clone, and we then averaged the values for the two clones sampled from each LTEE population at either 10,000 or 60,000 generations.

---

**Figure S1.**
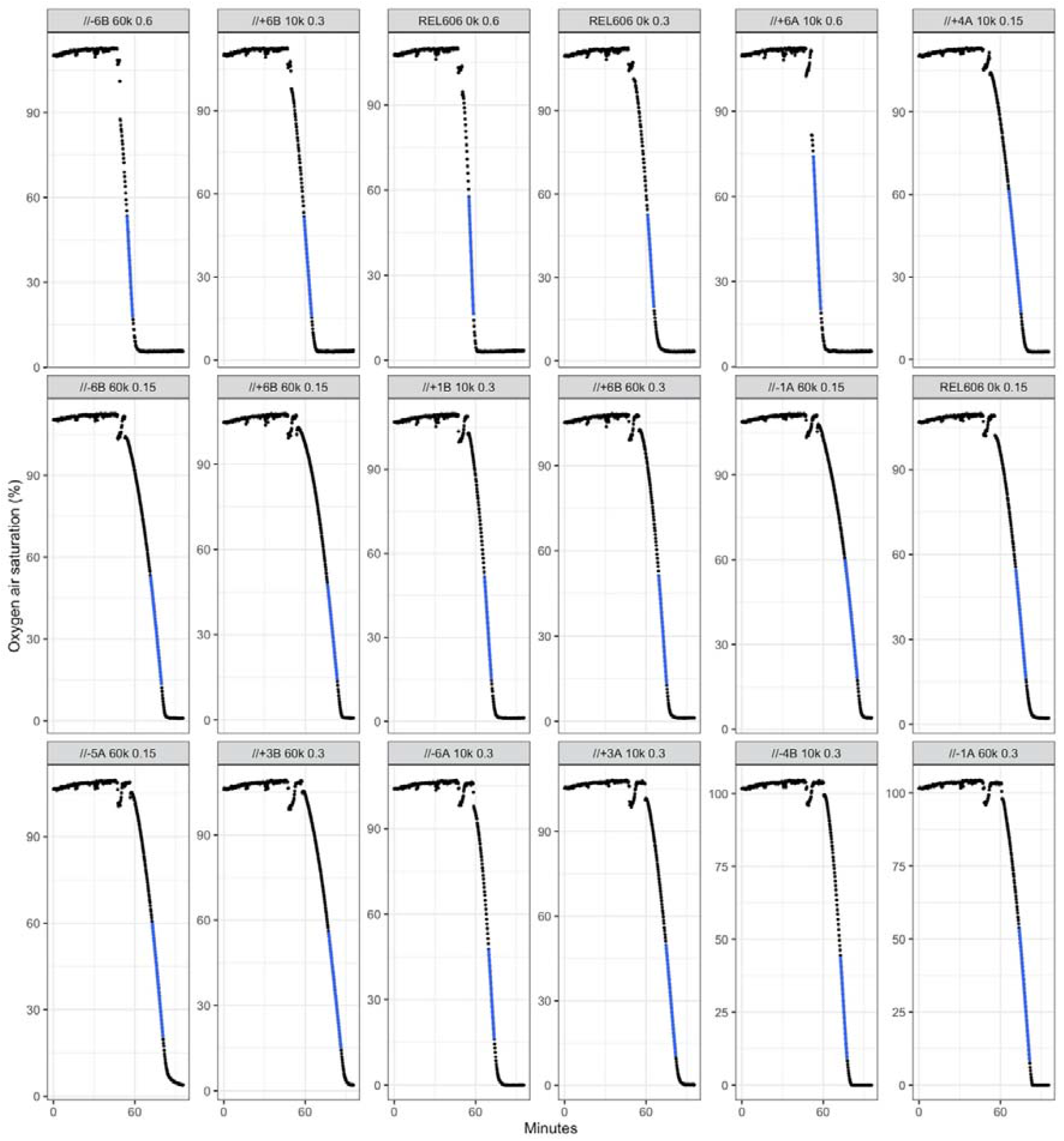
We monitored changes in oxygen saturation over time to estimate rates of energy use from the linear part of each time series (blue dots). Each panel shows a bacterial culture in which its initial biomass was standardized to a specific optical density and the oxygen was then monitored every 15 sec until it was largely or entirely depleted. Here we display the trajectories for 18 of the 192 time series in the full dataset (171 with cell cultures and 21 blanks).

**Figure S2.**
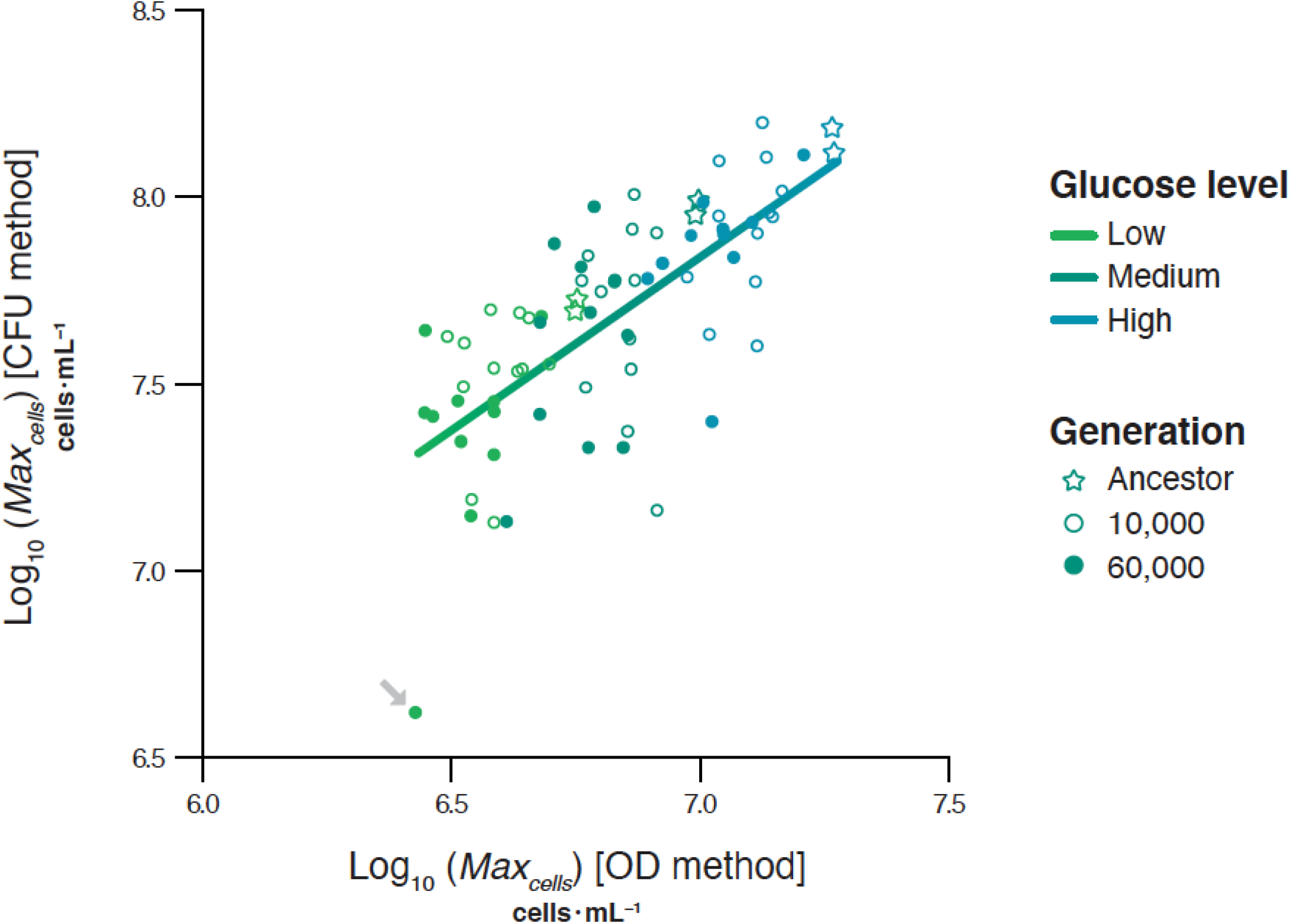
Relation between maximum cell density of each sample estimated based on the optical density of growing cultures and colony forming units at stationary phase. Growing cells are usually larger than cells in stationary phase, leading to lower estimates of the maximum cell density from the OD method and using this calibration. Each point shows the mean value for a sample (two clones from the same generation for the evolving populations), and symbols identify the different generations. The grey arrow indicates an apparent outlier (population Ara+3 at 60,000 generations), which had unusually low maximum densities using the CFU method. However, including or excluding this outlier has little effect on estimated scaling exponents (Table S1).

**Table S1.** *E. coli* ancestors and evolved clones used in this study.

| Clone ID | Population ID | Generation | Notes |
| --- | --- | --- | --- |
| REL606 | NA | 0 | Ara<sup>-</sup> ancestor |
| REL607 | NA | 0 | Ara<sup>+</sup> ancestor |
| REL4536A | NA | 10,000 | Clone A from Ara-1 population at 10,000 generations |
| REL4536B | NA | 10,000 | Clone B from Ara-1 population at 10,000 generations |
| REL4537A | NA | 10,000 | Clone A from Ara-2 population at 10,000 generations |
| REL4537B | NA | 10,000 | Clone B from Ara-2 population at 10,000 generations |
| REL4538A | NA | 10,000 | Clone A from Ara-3 population at 10,000 generations |
| ZDB429 | NA | 10,000 | Clone B from Ara-3 population at 10,000 generations |
| REL4539A | NA | 10,000 | Clone A from Ara-4 population at 10,000 generations |
| REL4539B | NA | 10,000 | Clone B from Ara-4 population at 10,000 generations |
| REL4540A | NA | 10,000 | Clone A from Ara-5 population at 10,000 generations |
| REL4540B | NA | 10,000 | Clone B from Ara-5 population at 10,000 generations |
| REL4541A | NA | 10,000 | Clone A from Ara-6 population at 10,000 generations |
| REL4541B | NA | 10,000 | Clone B from Ara-6 population at 10,000 generations |
| REL4530A | NA | 10,000 | Clone A from Ara+1 population at 10,000 generations |
| REL4530B | NA | 10,000 | Clone B from Ara+1 population at 10,000 generations |
| REL4531A | NA | 10,000 | Clone A from Ara+2 population at 10,000 generations |
| REL4531B | NA | 10,000 | Clone B from Ara+2 population at 10,000 generations |
| REL4532A | NA | 10,000 | Clone A from Ara+3 population at 10,000 generations |
| REL4532B | NA | 10,000 | Clone B from Ara+3 population at 10,000 generations |
| REL4533A | NA | 10,000 | Clone A from Ara+4 population at 10,000 generations |
| REL4533B | NA | 10,000 | Clone B from Ara+4 population at 10,000 generations |
| REL4534A | NA | 10,000 | Clone A from Ara+5 population at 10,000 generations |
| REL4534B | NA | 10,000 | Clone B from Ara+5 population at 10,000 generations |
| REL4535A | NA | 10,000 | Clone A from Ara+6 population at 10,000 generations |
| REL4535B | NA | 10,000 | Clone B from Ara+6 population at 10,000 generations |
| MJM 631 | REL 11678 | 60,000 | Clone A from Ara-1 population at 60,000 generations |
| MJM 632 | REL 11678 | 60,000 | Clone B from Ara-1 population at 60,000 generations |
| MJM 633 | REL 11679 | 60,000 | Clone A from Ara-2 population at 60,000 generations |
| MJM 634 | REL 11679 | 60,000 | Clone B from Ara-2 population at 60,000 generations |
| MJM 637 | REL 11681 | 60,000 | Clone A from Ara-4 population at 60,000 generations |
| MJM 638 | REL 11681 | 60,000 | Clone B from Ara-4 population at 60,000 generations |
| MJM 639 | REL 11682 | 60,000 | Clone A from Ara-5 population at 60,000 generations |
| MJM 640 | REL 11682 | 60,000 | Clone B from Ara-5 population at 60,000 generations |
| MJM 641 | REL 11763 | 60,000 | Clone A from Ara-6 population at 60,000 generations |
| MJM 642 | REL 11763 | 60,000 | Clone B from Ara-6 population at 60,000 generations |
| MJM 643 | REL 11765 | 60,000 | Clone A from Ara+1 population at 60,000 generations |
| MJM 644 | REL 11765 | 60,000 | Clone B from Ara+1 population at 60,000 generations |
| MJM 645 | REL 11686 | 60,000 | Clone A from Ara+2 population at 60,000 generations |
| MJM 646 | REL 11686 | 60,000 | Clone B from Ara+2 population at 60,000 generations |
| MJM 647 | REL 11687 | 60,000 | Clone A from Ara+3 population at 60,000 generations |
| MJM 648 | REL 11687 | 60,000 | Clone B from Ara+3 population at 60,000 generations |
| MJM 649 | REL 11688 | 60,000 | Clone A from Ara+4 population at 60,000 generations |
| MJM 650 | REL 11688 | 60,000 | Clone B from Ara+4 population at 60,000 generations |
| MJM 651 | REL 11723 | 60,000 | Clone A from Ara+5 population at 60,000 generations |
| MJM 652 | REL 11723 | 60,000 | Clone B from Ara+5 population at 60,000 generations |
| MJM 653 | REL 11724 | 60,000 | Clone A from Ara+6 population at 60,000 generations |
| MJM 654 | REL 11724 | 60,000 | Clone B from Ara+6 population at 60,000 generations |

**Table S2.** Modified version of Table 1 from the main text. As shown in Figure S2, one sample was an outlier in the calibration of maximum cell densities based on optical densities and colony forming units. Here we show that including or excluding that outlier has no meaningful effects on the observed scaling exponents.

| Parameter | Prediction<br>if $C = 0.11$ | Observed scaling<br>(including outlier) | Observed scaling<br>(excluding outlier) |
| --- | --- | --- | --- |
| $r$ | 0.27 | 0.27 | 0.27 |
| $Max_{cells}$ | -0.38 | -0.45 | -0.42 |
| $Max_{biovolume}$ | 0.62 | 0.55 | 0.58 |
| $Biovolume$<br>*productivity* | 0.89 | 0.81 | 0.85 |

## Literature Cited

1 Hatton, I. A. et al. The predator-prey power law: Biomass scaling across terrestrial and aquatic biomes. Science 349, (349), doi:10.1126/science.aac6284 (2015).

2 Malerba, M. E. & Marshall, D. J. Size-abundance rules? Evolution changes scaling relationships between size, metabolism and demography. Ecology Letters 22, 1407–1416, doi:10.1111/ele.13326 (2019).

3 Isaac, N. J. B., Storch, D. & Carbone, C. The paradox of energy equivalence. Global Ecology and Biogeography 22, 1–5, doi:10.1111/j.1466-8238.2012.00782.x (2013).

4 Hatton, I. A., Dobson, A. P., Storch, D., Galbraith, E. D. & Loreau, M. Linking scaling laws across eukaryotes. Proceedings of the National Academy of Sciences,USA 116, 21616–21622, doi:10.1073/pnas.1900492116 (2019).

5 Damuth, J. Population density and body size in mammals. Nature 290, 699–700, doi:10.1038/290699a0 (1981).

6 Savage, V. M., Gillooly, J. F., Brown, J. H., West, G. B. & Charnov, E. L. Effects of body size and temperature on population growth. American Naturalist 163, 429–441, doi:10.1086/381872 (2004).

7 Forster, J., Hirst, A. G. & Atkinson, D. Warming-induced reductions in body size are greater in aquatic than terrestrial species. Proceedings of the National Academy of Sciences, USA 109, 19310–19314, doi:10.1073/pnas.1210460109 (2012).

8 Audzijonyte, A. et al. Fish body sizes change with temperature but not all species shrink with warming. Nature Ecology & Evolution 4, 809–814, doi:10.1038/s41559-020-1171-0 (2020).

9 Gardner, J. L., Peters, A., Kearney, M. R., Joseph, L. & Heinsohn, R. Declining body size: a third universal response to warming? Trends in Ecology and Evolution 26, 285–291, doi:10.1016/j.tree.2011.03.005 (2011).

10 Audzijonyte, A., Kuparinen, A., Gorton, R. & Fulton, E. A. Ecological consequences of body size decline in harvested fish species: positive feedback loops in trophic interactions amplify human impact. Biology Letters 9, 20121103, doi:doi:10.1098/rsbl.2012.1103 (2013).

11 Bernhardt, J. R., Sunday, J. M. & O’Connor, M. I. Metabolic theory and the temperature-size rule explain the temperature dependence of population carrying capacity. American Naturalist 192, 687–697, doi:10.1086/700114 (2018).

12 Isaac, N. J. B., Carbone, C. & McGill, B. in Metabolic Ecology: A Scaling Approach (eds R. Sibly, J. H. Brown, & A. Kodric-Brown) 77–85 (2012).

13 Van M. Savage, James F. Gillooly, James H. Brown, Geoffrey B. West & Eric L. Charnov. Effects of body size and temperature on population growth. American Naturalist 163, 429–441, doi:10.1086/381872 (2004).

14 Hayward, A., Kolasa, J. & Stone, J. R. The scale-dependence of population density–body mass allometry: Statistical artefact or biological mechanism? Ecological Complexity 7, 115–124 (2010).

15 Kempes, C. P., Dutkiewicz, S. & and Follows, M. J. Growth, metabolic partitioning, and the size of microorganisms. Proceedings of the National Academy of Sciences, USA 109, 495–500 (2012).

16 Lenski, R. E., Rose, M. R., Simpson, S. C. & Tadler, S. C. Long-term experimental evolution in Escherichia coli. I. Adaptation and divergence during 2,000 generations. American Naturalist 138, 1315–1341, doi:10.1086/285289 (1991).

17 Good, B. H., McDonald, M. J., Barrick, J. E., Lenski, R. E. & Desai, M. M. The dynamics of molecular evolution over 60,000 generations. Nature 551, 45–50, doi:10.1038/nature24287 (2017).

18 Wiser, M. J., Ribeck, N. & Lenski, R. E. Long-term dynamics of adaptation in asexual populations. Science 342, 1364–1367, doi:10.1126/science.1243357 (2013).

19 Wortel, M. T., Noor, E., Ferris, M., Bruggeman, F. J. & Liebermeister, W. Metabolic enzyme cost explains variable trade-offs between microbial growth rate and yield. PLoS Computational Biology 14, 21, doi:10.1371/journal.pcbi.1006010 (2018).

20 Novak, M. et al. Experimental tests for an evolutionary trade-off between growth rate and yield in E. coli. American Naturalist 168, 242–251, doi:10.1086/506527 (2006).

21 Gallet, R. et al. The evolution of bacterial cell size: the internal diffusion-constraint hypothesis. ISME Journal 11, 1559–1568, doi:10.1038/ismej.2017.35 (2017).

22 Grant, N. A., Magid, A. A., Franklin, J., Dufour, Y. & Lenski, R. E. Changes in cell size and shape during 50,000 generations of experimental evolution with Escherichia coli. Journal of Bacteriology 203, e00469–20 doi: 10.1128/JB.00469-20 (2020).

23 Tenaillon, O. et al. Tempo and mode of genome evolution in a 50,000-generation experiment. Nature 536, 165–170, doi:10.1038/nature18959 (2016).

24 DeLong, J. P., Okie, J. G., Moses, M. E., Sibly, R. M. & Brown, J. H. Shifts in metabolic scaling, production, and efficiency across major evolutionary transitions of life. Proceedings of the National Academy of Sciences, USA 107, 12941–12945, doi:10.1073/pnas.1007783107 (2010).

25 García, F. C. et al. The allometry of the smallest: superlinear scaling of microbial metabolic rates in the Atlantic Ocean. ISME Journal 10, 1029–1036, doi:10.1038/ismej.2015.203 (2016).

26 Malerba, M. E., White, C. R. & Marshall, D. J. Phytoplankton size-scaling of net-energy flux across light and biomass gradients. Ecology 98, 3106–3115, doi:10.1002/ecy.2032 (2017).

27 Malerba, M. E., White, C. R. & Marshall, D. J. Eco-energetic consequences of evolutionary shifts in body size. Ecology Letters 21, 54–62, doi:10.1111/ele.12870 (2018).

28 Niklas, K. J. Plant Allometry: The Scaling of Form and Process. (2004).

29 Kozłowski, J., Konarzewski, M. & Gawelczyk, A. T. Cell size as a link between noncoding DNA and metabolic rate scaling. Proceedings of the National Academy of Sciences, USA 100, 14080–14085, doi:10.1073/pnas.2334605100 (2003).

30 Cooper, V. S. Long-term experimental evolution in Escherichia coli. X. Quantifying the fundamental and realized niche. BMC Evolutionary Biology 2, 12, doi:10.1186/1471-2148-2-12 (2002).

31 Philippe, N., Crozat, E., Lenski, R. E. & Schneider, D. Evolution of global regulatory networks during a long-term experiment with Escherichia coli. Bioessays 29, 846–860 (2007).

32 Favate, J. S., Liang, S., Yadavalli, S. S. & Shah, P. The landscape of transcriptional and translational changes over 22 years of bacterial adaptation. bioRxiv doi:10.1101/2021.01.12.426406 (2021).

33 Maddamsetti, R. et al. Core genes evolve rapidly in the long-term evolution experiment with Escherichia coli. Genome Biology and Evolution 9, 1072–1083, doi:10.1093/gbe/evx064 (2017).

34 Turner, C. B., Wade, B. D., Meyer, J. R., Sommerfeld, B. A. & Lenski, R. E. Evolution of organismal stoichiometry in a long-term experiment with Escherichia coli. Royal Society Open Science 4, 170497–170497, doi:10.1098/rsos.170497 (2017).

35 Mongold, J. A. & Lenski, R. E. Experimental rejection of a nonadaptive explanation for increased cell size in Escherichia coli. Journal of Bacteriology 178, 5333–5334, doi:10.1128/jb.178.17.5333-5334.1996 (1996).

36 Szenk, M., Dill, K. A. & de Graff, A. M. R. Why do fast-growing bacteria enter overflow metabolism? Testing the membrane real estate hypothesis. Cell Systems 5, 95–104, doi:10.1016/j.cels.2017.06.005 (2017).

37 Brown, J. H., Gillooly, J. F., Allen, A. P., Savage, V. M. & West, G. B. Toward a metabolic theory of ecology. Ecology 85, 1771–1789, doi:10.1890/03-9000 (2004).

38 Lynch, M. & Marinov, G. K. The bioenergetic costs of a gene. Proceedings of the National Academy of Sciences, USA 112, 15690–15695, doi:10.1073/pnas.1514974112 (2015).

39 Stouthamer, A. H. & Bettenhaussen, C. W. Continuous culture study of an APTase-negative mutant of Escherichia coli. Archives of Microbiology 113, 185–189, doi:10.1007/bf00492023 (1977).

40 Niklas, K. J. & Hammond, S. T. Biophysical effects on the scaling of plant growth, form, and ecology. Integrative and Comparative Biology 59, 1312–1323, doi:10.1093/icb/icz028 (2019).

41 Malerba, M. E., Ghedini, G. & Marshall, D. J. Genome size affects fitness in the eukaryotic alga Dunaliella tertiolecta. Current Biology, doi:10.1016/j.cub.2020.06.033 (2020).

42 Lenski, R. E. Experimental studies of pleiotropy and epistasis in Escherichia coli. I. Variation in competitive fitness among mutants resistant to virus T4. Evolution 42, 425–432 (1988).

43 Blount, Z. D., Borland, C. Z. & Lenski, R. E. Historical contingency and the evolution of a key innovation in an experimental population of Escherichia coli. Proceedings of the National Academy of Sciences, USA 105, 7899–7906 (2008).

44 Malerba, M. E., Palacios, M. M. & Marshall, D. J. Do larger individuals cope with resource fluctuations better? An artificial selection approach. Proceedings of the Royal Society of London B. 285, 1–9, doi:10.1098/rspb.2018.1347 (2018).

45 R Core Team. R: A Language and Environment for Statistical Computing. R Foundation for Statistical Computing, Vienna, Austria. URL http://www.R-project.org/ (2019).

46 Pinheiro, J., Bates, D., DebRoy, S., Sarkar, D. & R Core Team. nlme: linear and nonlinear mixed effects models. R package version 3.1-128, http://CRAN.R-project.org/package=nlme. (2016).

47 Bates, D., Mächler, M., Bolker, B. & Walker, S. Fitting linear mixed-effects models using lme4. Journal of Statistical Software 67, 1–48, doi:10.18637/jss.v067.i01 (2015).

48 Wickham, H. The split-apply-combine strategy for data analysis. Journal of Statistical Software 40, 1–29 (2011).

